# Genome-wide mapping of native co-localized G4s and R-loops in living cells

**DOI:** 10.1101/2024.06.03.597194

**Authors:** Ting Liu, Xing Shen, Yijia Ren, Hongyu Lu, Yu Liu, Chong Chen, Lin Yu, Zhihong Xue

**Affiliations:** Key Laboratory of Birth Defects and Related Disease of Women and Children of MOE, Department of Pediatrics, West China Second University Hospital, State Key Laboratory of Biotherapy and Collaborative Innovation Center of Biotherapy, Sichuan University, Chengdu, Sichuan, 610041, China; Department of Hematology and Institute of Hematology, State Key Laboratory of Biotherapy and Cancer Center, West China Hospital, Sichuan University, Chengdu, Sichuan, China

**Keywords:** G-quadruplex, G4, R-loop, Hemin, HepG4-seq, HBD-seq, Dhx9

## Abstract

The non-B DNA structures can act as dynamic functional genomic elements regulating gene expression. Among them, G4s and R-loops are two of the best studied. The interplay between R-loops and G4s are emerging in regulating DNA repair, replication and transcription. A comprehensive picture of native co-localized G4s and R-loops in living cells is currently lacking. Here, we describe the development of HepG4-seq and an optimized HBD-seq methods, which robustly capture native G4s and R-loops, respectively, in living cells. We successfully employed these methods to establish comprehensive maps of native co-localized G4s and R-loops in human HEK293 cells and mouse embryonic stem cells (mESCs). We discovered that co-localized G4s and R-loops are dynamically altered in a cell type-dependent manner and are largely localized at active promoters and enhancers of transcriptional active genes. We further demonstrated the helicase Dhx9 as a direct and major regulator that modulates the formation and resolution of co-localized G4s and R-loops. Depletion of Dhx9 impaired the self-renewal and differentiation capacities of mESCs by altering the transcription of co-localized G4s and R-loops - associated genes. Taken together, our work established that the endogenous co-localized G4s and R-loops are prevalently persisted in the regulatory regions of active genes and are involved in the transcriptional regulation of their linked genes, opening the door for exploring broader roles of co-localized G4s and R-loops in development and disease.

## Introduction

Genomic DNA can form various types of non-B secondary structures, including G-quadruplexes (G4s), R-loops, Z-DNA, i-motifs, Cruciform and others (Matos-Rodrigues et al., 2023). Among them, G4s and R-loops are two of the best studied. G4s are built by stacked guanine tetrads connected via Hoogsteen hydrogen bonds and can be formed by intra- or inter-molecular folding of the tetramers (Panyutin et al., 1989; Sen and Gilbert, 1988; Sundquist and Klug, 1989; Williamson et al., 1989). R-loops are three-stranded structures containing a DNA-RNA hybrid and a displaced single-stranded DNA (Aguilera and Garcia-Muse, 2012; Xu and Clayton, 1996). Both G4s and R-loops are involved in key biological processes, including transcription, replication, genomic instability, class switch recombination in B cells, DNA damage and repair, and telomere maintenance (Garcia-Muse and Aguilera, 2019; Robert Hänsel-Hertsch, 2017; Varshney et al., 2020; Yang et al., 2023).

R-loops appear to have a strong sequence preference with high G/C ratios (Ginno et al., 2013; Ginno et al., 2012). Reports about the interplay between R-loops and G4s are emerging. Specific G4 ligands stabilized G4s and simultaneously increase R-loop levels within minutes in human cancer cells, which finally induced DNA damage (De Magis et al., 2019). Reactive oxygen species (ROS) have been reported to induce G4 and R-loop formation at transcriptionally active sites, and their inter-regulation is essential for the DNA repair (Tan et al., 2020). In a reconstituted eukaryotic DNA replication system, the interplay of R-loops and G4s was shown to impact replication fork progression by inducing fork stalling (Kumar et al., 2021). Single-molecule fluorescence studies showed the existence of a positive feedback mechanism of G4 and R-loop formation during transcription, where the transcription-induced R-loop precedes and facilitates G4 formation in the non-template strand, and in turn G4 promotes the R-loop formation in the following rounds of transcription (Lee et al., 2020; Lim and Hohng, 2020). Wulfridge et al. reported that the architectural protein CCCTC binding factor (CTCF)-bound sites are enriched for R-loops and G4s which facilitate CTCF binding to promote chromatin looping interactions (Wulfridge et al., 2023).

Detection of G4s and R-loops have been largely based on the use of a single-chain variable fragment (scFv) BG4 for G4s and a monoclonal antibody S9.6 for R-loops. In recent years, these two antibodies have been coupled with deep sequencing to genome-widely detect G4s and R-loops (Galli et al., 2022; Ginno et al., 2012; Hänsel-Hertsch et al., 2016; Jiang et al., 2023; Lyu et al., 2021). Using BG4 and S9.6 -based CUT & Tag, G4s and R-loops showed high degree of co-occurrence in mESCs (Lyu et al., 2021). However, given that a group of helicases, RNA-binding factors, endonucleases and DNA topoisomerases cooperate to actively dissolve G4s and R-loops restoring B-formed DNA duplexes (Robert Hänsel-Hertsch, 2017; Varshney et al., 2020; Yang et al., 2023), a steady state equilibrium is generally set at low levels in living cells under physiological conditions (Miglietta et al., 2020) and thus addition of high affinity antibodies may pull the equilibrium towards folded states. Additionally, the specificity of the S9.6 antibody on R-loops has been questioned recently for accurate quantification and mapping of R-loops (Hartono et al., 2018; Konig et al., 2017; Phillips et al., 2013).

To understand the co-localized G4s and R-loops in living cells under physiological conditions, we sought to develop an *in vivo* strategy for G4 profiling based on the G4-hemin complex-induced proximal labeling and R-loop profiling based on the N-terminal hybrid-binding domain (HBD) of RNase H1. Recent studies showed that G4s could tightly form a complex with the cellular cofactor hemin both *in vitro* and in living cells, where hemin binds by end-stacking on the terminal G-quartets of G4s without affecting the folding of G4s (Gray et al., 2019; Stadlbauer et al., 2021). The G4-hemin complex has been shown to act as a peroxidase to catalyze oxidation reactions in the presence of hydrogen peroxide (H_2_O_2_) (Cheng et al., 2009; Lat et al., 2020; Yang et al., 2011). The H_2_O_2_-activated G4-hemin complex oxidizes the biotin tyramide to phenoxyl redicals that covalently conjugate biotin to G4 itself and its proximal DNA within 10 nm (equivalent to approximately 31 bp) *in vitro* and *in vivo* (Einarson and Sen, 2017; Lat et al., 2020). Here, we have utilized the G4-hemin-mediated proximal biotinylation rection to develop a new method HepG4-seq (for high throughput **seq**uencing of **he**min-induced **p**roximal labelled **G4**s) to map the genomic native G4s under physiological conditions. The HBD domain of RNase H1 mediates the specific recognition of DNA/RNA hybrid in a sequence-independent manner, which is a gold standard for R-loop recognition in the cell (Nowotny et al., 2008). The catalytically inactive RNase H1 or its HBD domain -based methods have been successfully used to identify genome-wide native R-loops (Chen et al., 2017; Wang et al., 2021). We have adapted the “GST-His6-2xHBD”-mediated CUT&Tag protocol (Wang et al., 2021) to develop the HBD-seq protocol in this study.

We have combined the HepG4-seq and HBD-seq to profile the genome-wide native co-localized G4s and R-loops with high signal-to-noise ratios in HEK293 cells and mouse embryonic stem cells (mESCs). We observed that the co-localized G4s and R-loops exhibit cell type-dependent distributions and are largely localized at active promoters and enhancers of transcriptionally active genes. We further showed that ∼70% of the co-localized G4s and R-loops in mESCs were directly bound by the helicase Dhx9 and that depletion of Dhx9 significantly altered the levels of ∼6200 co-localized G4s and R-loops bound by Dhx9. Furthermore, depletion of Dhx9 was shown to impair the self-renewal and differentiation capacities of mESCs by altering the transcription of co-localized G4s and R-loops -associated genes.

## Results

### Mapping of the native DNA G4 through the G4-hemin-mediated proximal biotinylation

The DNA G4-hemin complex could act as a mimic peroxidase to oxidize the biotin tyramide to phenoxyl radicals that can covalently conjugate biotin to G4 itself and its proximal DNA within ∼30bp in the presence of H_2_O_2_ (Cheng et al., 2009; Einarson and Sen, 2017; Lat et al., 2020). However, the efficiency of peroxidase-mediated biotinylation on DNA is limited using the substrate biotin tyramide (Zhou et al., 2019). Recently, biotin aniline (Bio-An) has been shown to have superior labeling efficiency on DNA than biotin tyramide, when catalyzed by the engineered peroxidase APEX2 (Zhou et al., 2019). The free heme concentration in normal human erythrocytes is 21 ± 2 µM (Aich et al., 2015). To explore the G4-hemin-mediated biotinylation in the living cells, we treated HEK293 cells with 25 µM hemin and 500 µM Bio-An for 2 hours prior to activation with 1mM H_2_O_2_ for 1 minute, and then quenched the labeling reaction and performed the immunofluorescence staining using Alexa Fluor 647 conjugated recombinant streptavidin (Strep-647) that specifically recognizes biotin. As shown in Fig. 1A, cells treated with hemin and Bio- An exhibited a robust fluorescence signal, while the absence of either hemin or Bio- An almost completely abolished the biotinylation signals, suggesting a specific and active biotinylation activity. To understand whether addition of hemin disturbs the formation of G4s, we performed the BG4 CUT&Tag-seq using the recombinant BG4 and Tn5 on HEK293 cells treated with and without 25 µM hemin (Supplementary Fig. 1A, B). The heatmap and profile plot analysis showed similar BG4 CUT&Tag signals between the hemin-treated and control samples (Supplementary Fig. 1B). There were only 174 BG4 CUT&Tag peaks with significantly differential signals between the hemin-treated and control samples (Supplementary Fig. 1C). These data suggest that the hemin treatment condition we used does not significantly affect G4 folding. Therefore, hemin-induced proximal biotinylation of G4s could be utilized to mark the native G4s in living cells.

**Fig. 1.**
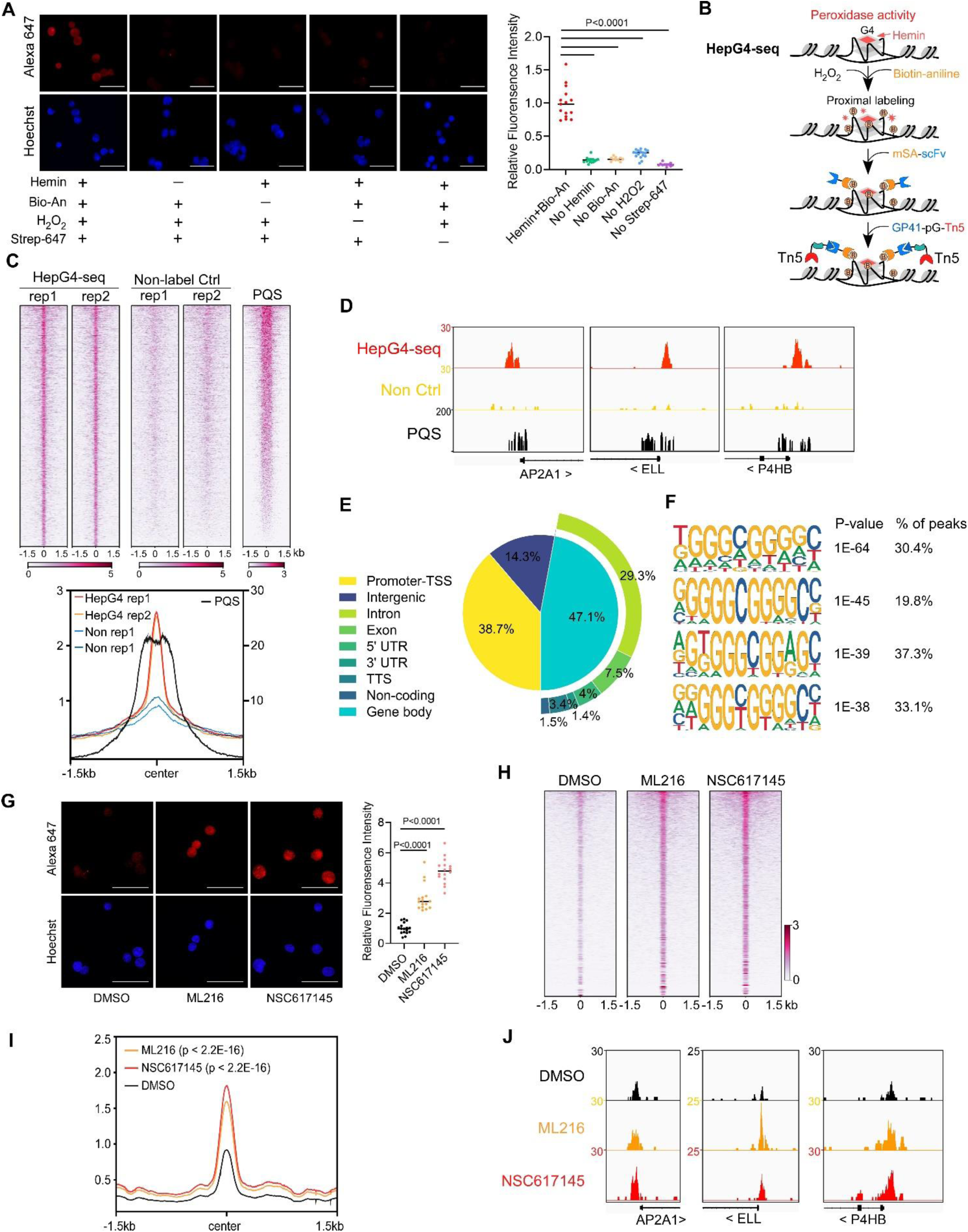
Mapping of the native DNA G4 through the G4-hemin-mediated proximal biotinylation. **A**, Immunofluorescence staining of the HEK293 cells treated with indicated conditions using the Alexa Fluor 647 labelled recombinant streptavidin (Strep-647). Nuclei were stained with the Hoechst33342. Scale bar, 50 µm. Bio-An, Biotin-aniline; H_2_O_2_, Hydrogen peroxide. The quantified relative fluorescence intensities were shown in the right panel. **B**, Schematic of the HepG4-seq procedure. SA-scFv, the recombinant fusion protein of mSA and anti-GP41 Single Chain Fragment Variable (scFv); GP41-pG-Tn5, the recombinant fusion protein of the GP41 tag, protein G and Tn5. **C**, Top: Heatmap showing the signal of BG4-seq, HepG4-seq and maxScores of PQS ±1.5kb around the center of peaks identified by HepG4-seq in HEK293 cells. Color scales represent the density of the signals. Bottom: Profile plot showing the average signal of HepG4-seq reads ±1.5kb around the center of peaks and the average maxScores of PQS calculated by pqsfinder at the same positions. HepG4 rep1/rep2, two biologically independent HepG4-seq replicates in HEK293 cells; Non rep1/rep2, two biologically independent non-label negative control replicates in HEK293 cells. **D**, Representative genome browser tracks showing HepG4-seq (red), non-label negative control of HepG4-seq (yellow), and PQS (black) signals in HEK293 cells along the indicated genomic loci. **E**, Distribution of HepG4-seq signals in HEK293 cells in different gene features. **F**, The top enriched motifs on the HepG4-seq peaks in HEK293 cells. **G**, Immunofluorescence staining of the HEK293 cells treated with DMSO, ML216 (25 µM), or NSC617145 (3 µM) using the Alexa Fluor 647 labelled recombinant streptavidin. Nuclei were stained with the Hoechst33342. Scale bar, 50 µm. The quantified relative fluorescence intensities were shown in the right panel. **H**, Heatmap showing the HepG4-seq signals ±1.5kb around the center of peaks identified in HEK293 cells treated with DMSO, ML216 (25 µM), or NSC617145 (3 µM). Color scales represent the density of the signals. **I**, Profile plot showing the average signal of HepG4-seq reads ±1.5kb around the center of peaks identified in HEK293 cells treated with DMSO, ML216 (25 µM), or NSC617145 (3 µM). The p values (ML216 v.s. DMSO; NSC617145 v.s. DMSO) were calculated using the Mann-Whitney test. **J**, Representative genome browser tracks showing the HepG4-seq signals in HEK293 cells treated with DMSO, ML216 (25 µM), or NSC617145 (3 µM) along the indicated genomic loci.

The recombinant streptavidin monomer (mSA) combines the streptavidin and rhizavidin sequences to achieve specific monovalent detection of biotin or biotinylated molecules with a high affinity (Kd= 2.8 nM) (Lim et al., 2013). The Moon-tag system consists of a 15-amino acid peptide GP41-tag and a 123-amino acid anti-GP41-tag nanobody with an affinity of ∼30 nM *in vitro* (Boersma et al., 2019). We expressed and purified the recombinant mSA fused with the anti-GP41 nanobody (mSA-scFv), and the recombinant Tn5 fused with the GP41-tag and protein G (GP41-pG-Tn5) from *E. coli* (Supplementary Fig. 1A). To map the hemin-induced biotinylated G4s, we developed a new method HepG4-seq (Fig. 1B), where mSA-scFv recognizes the biotinylated G4s and recruits the transposase Tn5 to achieve “Cleavage Under Targets and Tagmentation (CUT&Tag)”. Deep sequencing analysis of the biotinylated G4s fragments identified 6,799 consensus peaks from two independent biological repeats in HEK293 cells, where the signals were dramatically diminished in HEK293 cells without treatment with hemin and Bio-An, suggesting the specificity of HepG4-seq (Fig. 1C, Supplementary Table 1). Several representative HepG4s-seq-identified G4 peaks were shown in Fig. 1D. Genomic distribution analysis showed that G4s are mainly localized in promoters (38.7%) and gene bodies (47.1%) (Fig. 1E).

We also evaluated the HepG4-seq-identified peaks using a G4-forming sequences (PQS) predication tool pqsfinder which has been shown to have 96% accuracy on ∼400 known and experimentally observed G4 structures (Hon et al., 2017). The peaks identified by HepG4-seq overlap quite well with the center of pqsfinder maxScores that report the PQS quality (Fig. 1C-D). The motif enrichment analysis by HOMER (Heinz et al., 2010) revealed a high prevalence of G-rich sequences in HepG4-seq peaks (Fig. 1F). All above further validated the specificity of HepG4-seq in capturing G4s.

### Induction of DNA G4s by inhibiting G4 resolving helicase

The RecQ-like helicases Bloom syndrome protein (BLM) and Werner syndrome ATP-dependent helicase (WRN) are the first recognized and the best characterized DNA G4-resolving mammalian helicases (Fry and Loeb, 1999; Mendoza et al., 2016; Mohaghegh et al., 2001). The small molecules ML216 and NSC617145 are selective and cell permeable inhibitors of BLM and WRN, respectively, by inhibiting their ATPase activity (Aggarwal et al., 2013a; Aggarwal et al., 2013b; Nguyen et al., 2013). To investigate the effect of BLM or WRN inhibition on native G4s, we treated HEK293 cells with ML216 or NSC617145 for 16 hours and then labeled cellular G4s by hemin-G4-induced biotinylation in living cells. Immunofluorescence staining of the treated cells using Strep-647 showed that the treatment of ML216 or NSC617145 remarkably elevated signals of native G4s (Fig. 1G). Furthermore, we performed HepG4-seq on HEK293 cells treated with ML216 or NSC617145. Notably, HepG4-seq identified 77,003 peaks from ML216- or NSC617145-treated HEK293 cells, and ∼ 70,000 new G4 peaks were induced by inhibition of BLM or WRN (Supplementary Table 1). The signals of G4s detected by HepG4-seq were significantly increased after inhibiting BLM or WRN (Fig. 1H-I) (Mann-Whitney test, p < 2.2E-16). Representative G4s peaks are shown in Fig. 1J. In addition, the top enriched motifs in the 70000 extra HepG4-seq peaks are G-riched (Supplementary Fig. 1E). Taken together, these data suggest that HepG4-seq is able to efficiently detect dynamic native G4s.

### Mapping of native co-localized G4s and R-loops in HEK293 cells

The HBD domain of RNase H1 has been demonstrated as a DNA/RNA hybrid recognition sensor and applied to identify genome-wide native DNA/RNA hybrids using the recombinant GST-His6-2xHBD coupling with Tn5-based CUT&Tag (Nowotny et al., 2008; Wang et al., 2021). Given that the GST-fusion proteins are prone to form variable high molecular-weight aggregates and these aggregates often undermine the reliability of the fusion proteins (Deceglie et al., 2014; Ki and Pack, 2020), we produced the recombinant two copies of HBDs fused with EGFP and V5-tag (HBD-V5) (Supplementary Fig. 1A) and used the anti-V5 tag antibody instead of the anti-His tag antibody for the CUT&Tag-seq. We call this modified protocol as HBD-seq for mapping the native R-loops in cells (Fig. 2A). We performed the HBD-seq on HEK293 cells, and revealed 42,488 consensus native R-loops peaks with a high signal-to-noise ratio while the HBD-seq signals were dramatically diminished in HEK293 cells treated with the RNases prior to HBD-seq (Fig. 2B), suggesting the specificity of HBD-seq in detecting native R-loops.

**Fig. 2.**
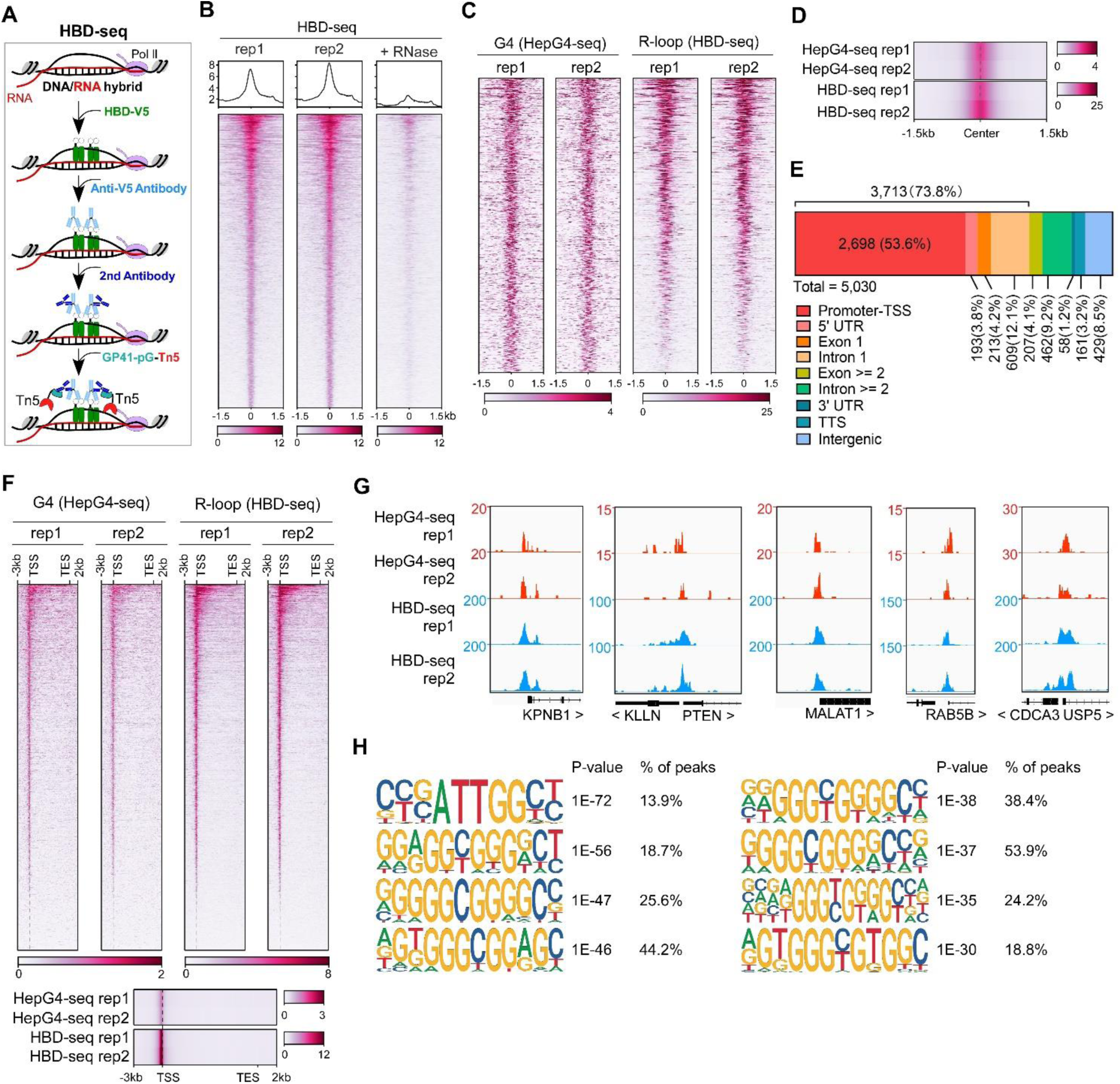
Mapping of the co-localized G4 and R-loop in the HEK 293 cells by combining the HepG4-seq and HBD-seq. **A**, Schematic of the HBD-seq procedure. HBD-V5, the recombinant fusion protein of the N-terminal hybrid-binding domain (HBD) of RNase H1 and V5 tag; GP41-pG-Tn5, the recombinant fusion protein of the GP41 tag, protein G and Tn5. **B**, Heatmap showing the signal of HBD-seq reads ±1.5kb around the center of peaks in HEK293 cells. Two biologically independent replicates are shown. “+ RNase” represents the treatment of RNase A and RNase H prior to the HBD-seq. Color scales represent the density of the signals. **C**, Heatmap showing the signal of HepG4-seq and HBD-seq reads ±1.5kb around the center of co-localized G4 and R-loop peaks in HEK293 cells. Color scales represent the density of the signals. **D**, Profile plot showing the average signal of HepG4-seq and HBD-seq reads ±1.5kb around the center of co-localized G4 and R-loop peaks in HEK293 cells. The plot is visualized using the heatmap. Color scales represent the density of the signals. **E**, Distribution of the co-localized G4s and R-loops in HEK293 cells in different gene features. **F**, Top: Heatmap showing the signal of HepG4-seq and HBD-seq reads of the co-localized G4s and R-loops in HEK293 cells along the gene body, 3kb upstream of transcription start site (TSS) and 2kb downstream of transcription end site (TES). Bottom: Profile plot showing the average signal of HepG4-seq and HBD-seq reads of the co-localized G4s and R-loops in HEK293 cells along the indicated gene features. The plot is visualized using the heatmap. Color scales represent the density of the signals. **G**, Representative genome browser tracks showing the HepG4-seq and HBD-seq signals of the co-localized G4s and R-loops in HEK293 cells along the indicated genomic loci. **H**, The top enriched motifs of the co-localized G4s and R-loops in HEK293 cells.

We then analyzed the regions co-occupied by both HepG4-seq-identified G4s and HBD-seq-identified R-loops, and revealed 5030 native co-localized peaks in HEK293 cells, ranging in size from 100 bp to ∼1.5 kb (Fig. 2C-D, Supplementary Fig. 1D). 73.8% of these co-localized peaks are localized at promoters, 5’UTR, exon1 and intron 1 (Fig. 2E). When we performed a metagene analysis of these co-localized peaks, a distinct peak was detected around the transcription start site (TSS) (Fig. 2F). Representative co-localized peaks are shown in the Fig. 2G. The motifs enrichment analysis by HOMER (Heinz et al., 2010) showed that G-rich sequences are highly enriched in the co-localized peaks (Fig. 2H).

### The co-localized G4s and R-loops-mediated transcriptional regulation in HEK293 cells

The predominant distribution of co-localized peaks around TSS implies that they may participate in transcriptional regulation of their associated genes. RNA-seq analysis revealed that the RNA levels of co-localized G4s and R-loops-associated genes are significantly higher than all genes, G4s or R-loops-associated genes with the Mann-Whitney test p < 2.2E-16 (Fig. 3A). Different from G4s and R-loops, the co-localized G4s and R-loops are mainly localized within 1 kb of the TSS of transcriptionally active genes (∼60% peaks with FPKM >=5) (Fig. 3B). To investigate the transcriptional regulation of co-localized G4s and R-loops in living cells, we performed the RNA-seq on HEK293 cells treated with and without ML216 or NSC617145 and then analyzed the differential gene expression using DESeq2 (Love et al., 2014). As a result, hundreds of genes were linked to co-localized G4s and R-loops with increased G4 signals (at least 1.5 foldchange) and at the same time exhibited significant changes in expression levels upon inhibition of BLM or WRN in HEK293 cells (Fig. 3C), suggesting that co-localized G4s and R-loops could regulate the transcription of their associated genes. Distribution analysis showed that these differential co-localized G4s and R-loops are mainly localized in the promoter-TSS (Fig. 3D). Among the differential genes, 125 genes were co-regulated by both ML216 and NSC617145 (Fig. 3E), suggesting that BLM and WRN could co-regulate the transcription of genes by resolving G4s. Gene ontology (GO) analysis showed that co-localized G4s & R-loops-regulated genes in HEK293 cells are mainly involved in cell cycle regulation, DNA/mRNA metabolic regulation, DNA damage response, chromatin binding, kinase binding, cell-substrate junction, et al (Fig. 3F).

**Fig 3.**
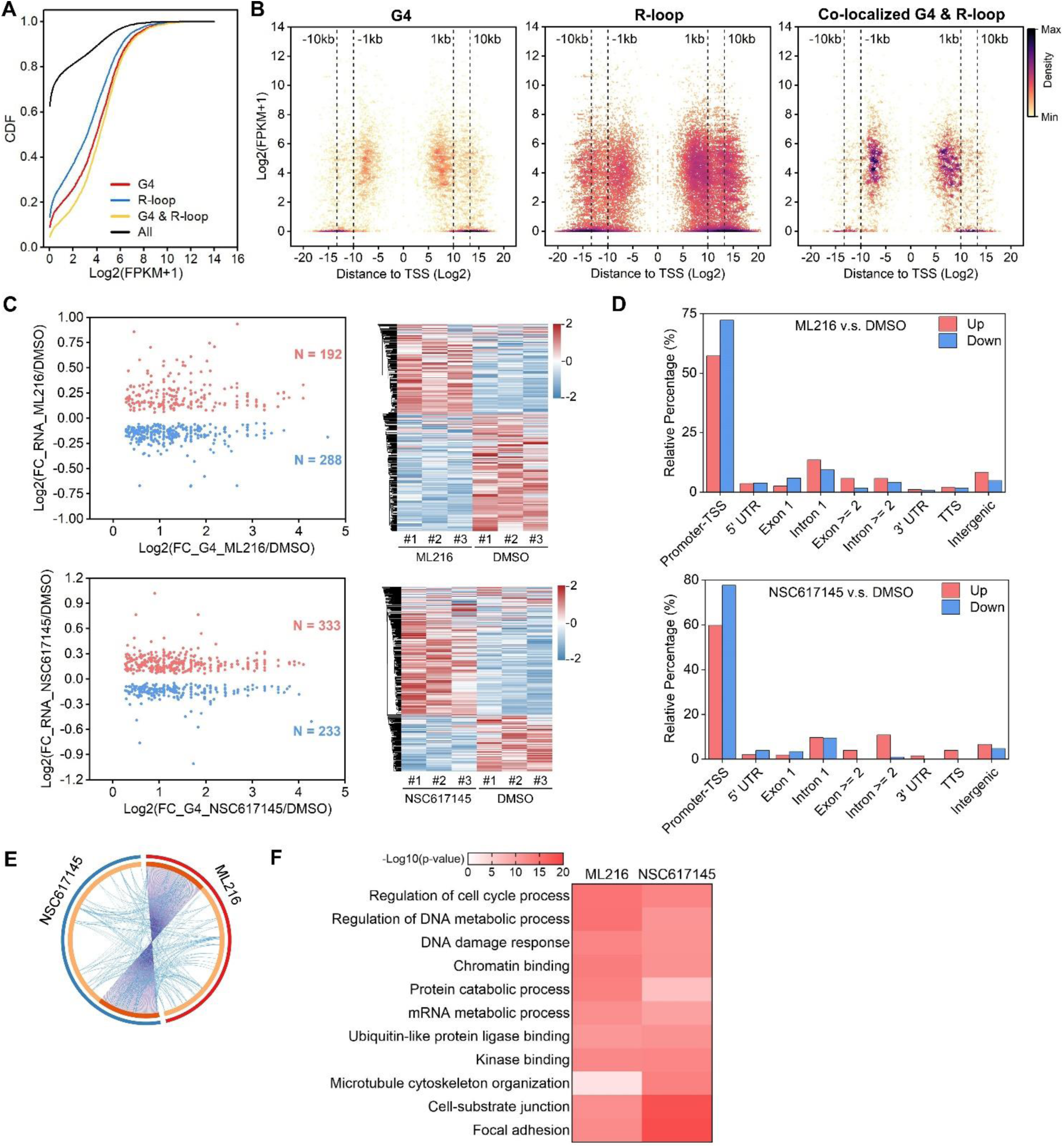
The co-localized G4s and R-loops-mediated transcriptional regulation in HEK293 cells. **A**, Cumulative distribution plot showing comparisons of FPKMs of G4, R-loop, co-localized G4 & R-loop associated genes and all genes in HEK293 cells. **B**, Scatter plot showing the distributions of FPKMs of G4, R-loop, co-localized G4 & R-loop-associated genes versus the distances of G4, R-loop, co-localized G4 & R-loop to nearest TSS. The distance is in bp. Color scales represent the density of dots. **C**, Left: Scatter plot showing the distributions of foldchanges (FCs) of RNA levels of co-localized G4 & R-loop -associated genes (p value < 0.05) versus FCs of G4 signals of co-localized G4s & R-loops (FC >=1.5) after treatment with indicated inhibitors of G4 resolving helicases BLM or WRN. The number of genes were labelled on the plot. Right: Heatmap showing differential expression levels of co-localized G4 & R-loop-associated genes after treatment with indicated G4 inhibitors. RNA-seq data are from three biologically independent repeats. Color scales represent the normalized expression levels. **D**, Distributions of co-localized G4 s& R-loops expressions of which-associated genes were significantly up- or down-regulated after treatment with indicated G4 inhibitors across different gene features. **E**, Circos plot showing the overlap co-localized G4 & R-loop-associated genes differentially expressed after the treatment of ML216 or NSC617145 in HEK293 cells. Purple lines link the same gene that are shared by multiple groups. Blue lines link the genes, although different, fall under the same ontology term. Dark orange color of the inside arc represents the genes that are shared by multiple groups and light orange color of the inside arc represents genes that are unique to that group. **F**, Heatmap showing the GO-based enrichment terms of co-localized G4 & R-loop-associated genes differentially expressed after the treatment of ML216 or NSC617145 in HEK293 cells. The heatmap cells are colored by their p-values.

### Mapping of native co-localized G4s and R-loops in mESCs

Mouse embryonic stem cells (mESCs) are pluripotent stem cells that could differentiate into various types of cells of three germ lineages (Murry and Keller, 2008; Young, 2011). To understand the regulatory roles of co-localized G4s and R-loops in mESCs, we performed the HepG4-seq and HBD-seq on mESCs and finally uncovered 68,482 native overlapping peaks in mESCs, ranging in size from 100 bp to ∼2 kb (Fig. 4A-D, Supplementary Fig. 2A, Supplementary Table 2). Notably, unlike HEK293 cells, large number of native G4s (95,128) were identified by HepG4-seq in mESCs (Fig. 4A, Supplementary Table 2), which well overlap with the PQS predicted by pqsfinder (Fig. 4B), suggesting that native G4s exhibit obvious cell type-specific distribution.

**Fig. 4.**
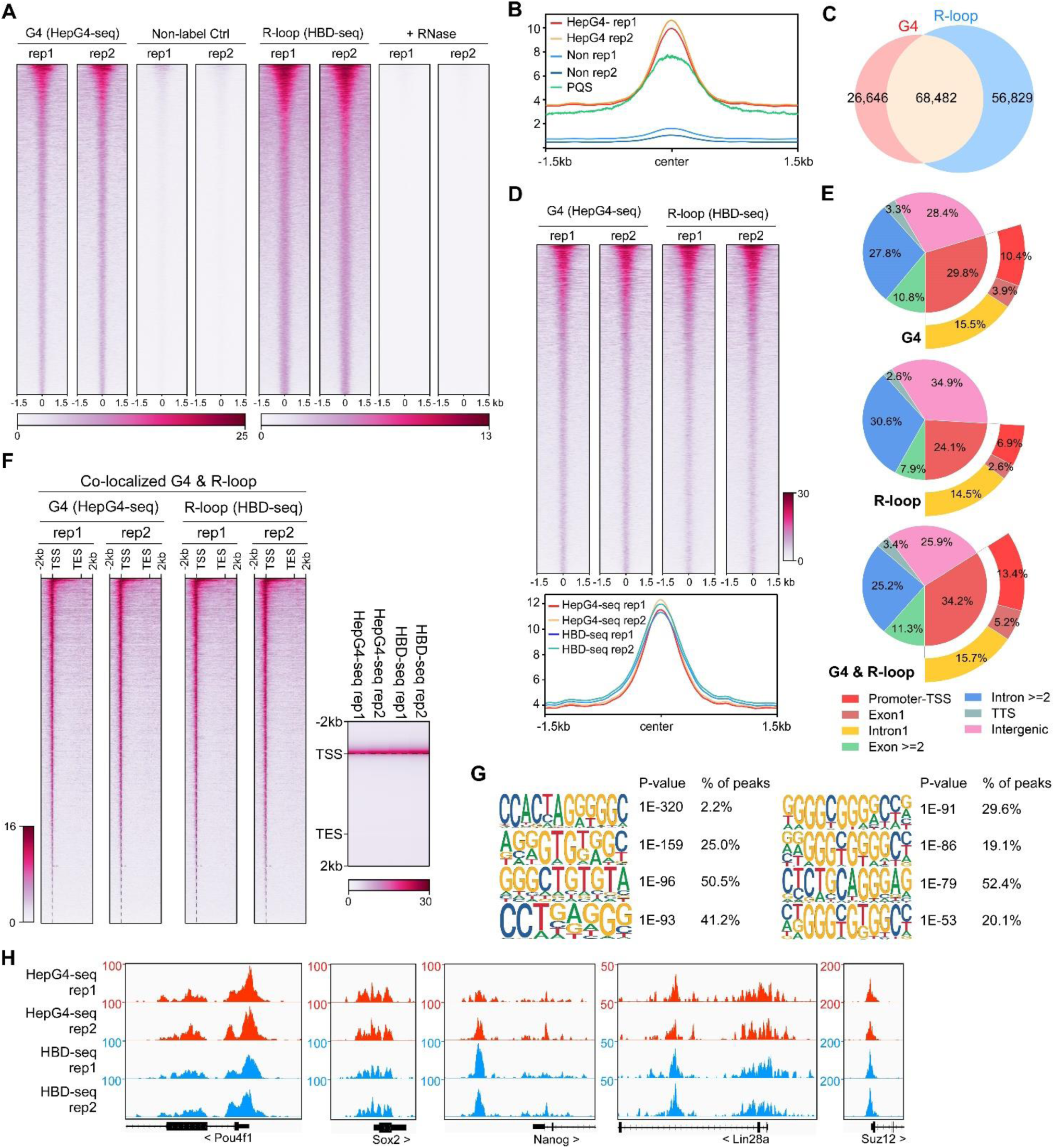
Mapping of the co-localized G4s and R-loops in the mouse embryonic stem cells. **A**, Heatmap showing the signal of HepG4-seq and HBD-seq ±1.5kb around the center of peaks in mESCs. Two biologically independent replicates are shown. Two biologically independent non-label replicates were the negative controls for HepG4-seq. Two biologically independent replicates with treatment of RNase A and RNase H prior to HBD-seq were the negative controls for HBD-seq. Color scales represent the density of the signals. **B**, Profile plot showing the average signal of HepG4-seq reads ±1.5kb around the center of peaks and the average maxScores of PQS calculated by pqsfinder at the same positions. HepG4 rep1/rep2, two biologically independent HepG4-seq replicates in mESCs; Non rep1/rep2, two biologically independent non-label negative control replicates in mESCs. **C**, Venn diagram comparing the DNA G4 and R-loop in mESCs. **D**, Top: Heatmap showing the signal of HepG4-seq and HBD-seq ±1.5kb around the center of co-localized G4s & R-loops in mESCs. Two biologically independent replicates are shown. Color scales represent the density of the signals. Bottom: Profile plot showing the average signal of HepG4-seq and HBD-seq reads ±1.5kb around the center of co-localized G4s & R-loops in mESCs. **E**, Distribution of G4s, R-loops, and co-localized G4s & R-loops signals in mESCs in different gene features. **F**, Left: Heatmap showing the signal of HepG4-seq and HBD-seq reads of the co-localized G4s & R-loops in mESCs along the gene body, 2kb upstream of TSS and 2kb downstream of TES. Right: Profile plot showing the average signal of HepG4-seq and HBD-seq reads of the co-localized G4s & R-loops in mESCs along the indicated gene features. The plot is visualized using the heatmap. Color scales represent the density of the signals. **G**, The top enriched motifs of the co-localized G4s & R-loops in mESCs. **H**, Representative genome browser tracks showing the HepG4-seq and HBD-seq signals of the co-localized G4s & R-loops in mESCs along the indicated genomic loci.

For the genomic distribution of co-localized G4s and R-loops, unlike HEK293 cells, only 34.2% peaks are localized in promoters, exon1 and intron1 while 25.9% peaks in intergenic regions (Fig. 4E). The distinct number and localization feature of co-localized G4s and R-loops in HEK293 cells and mESCs shows the cell type-specific distribution. The metagene analysis of overlapping peaks exhibited a distinct peak around TSS (Fig. 4F), suggesting the potential of transcriptional regulation. The motifs enrichment analysis found that G-rich sequences are highly enriched in these overlapping peaks, similar to those in HEK293 cells (Fig. 4G). Representative peaks were found in several key regulatory genes of mESCs (Fig. 4H).

### Characterization of native co-localized G4s and R-loops in mESCs

Similar to HEK293 cells, the RNA levels of co-localized G4s and R-loops-associated genes were seen to be significantly higher in mESCs (Mann-Whitney test p < 2.2E-16) (Fig. 5A). However, unlike in HEK293 cells (Fig. 3B), the overlapping peaks in mESCs are mainly localized in the proximal promoters (1kb from TSS, 9063 peaks with FPKM >=5) and the region 5-50 kb from the TSS of transcriptionally active genes (32690 peaks with FPKM >=5) (Fig. 5B), suggesting that co-localized G4s and R-loops are possibly distributed in active promoters or enhancers. To test this idea, we analyzed the co-localization of G4 (HepG4-seq), R-loop (HBD-seq), ChIP-seq signals of multiple chromatin markers and RNA polymerase II with the phosphorylated serine 5 at its CTD domain (RNAP) that marks the transcriptionally initiated RNA Polymerase II (Hsin and Manley, 2012). As a result, co-localized G4s and R-loops were observed to well overlap with active chromatin markers (H3K4me3, H3K27ac, H3K36me3, H3K4me1) and RNAP but not the repressed chromatin marker H3K27me3 (Fig. 5C).

**Fig. 5.**
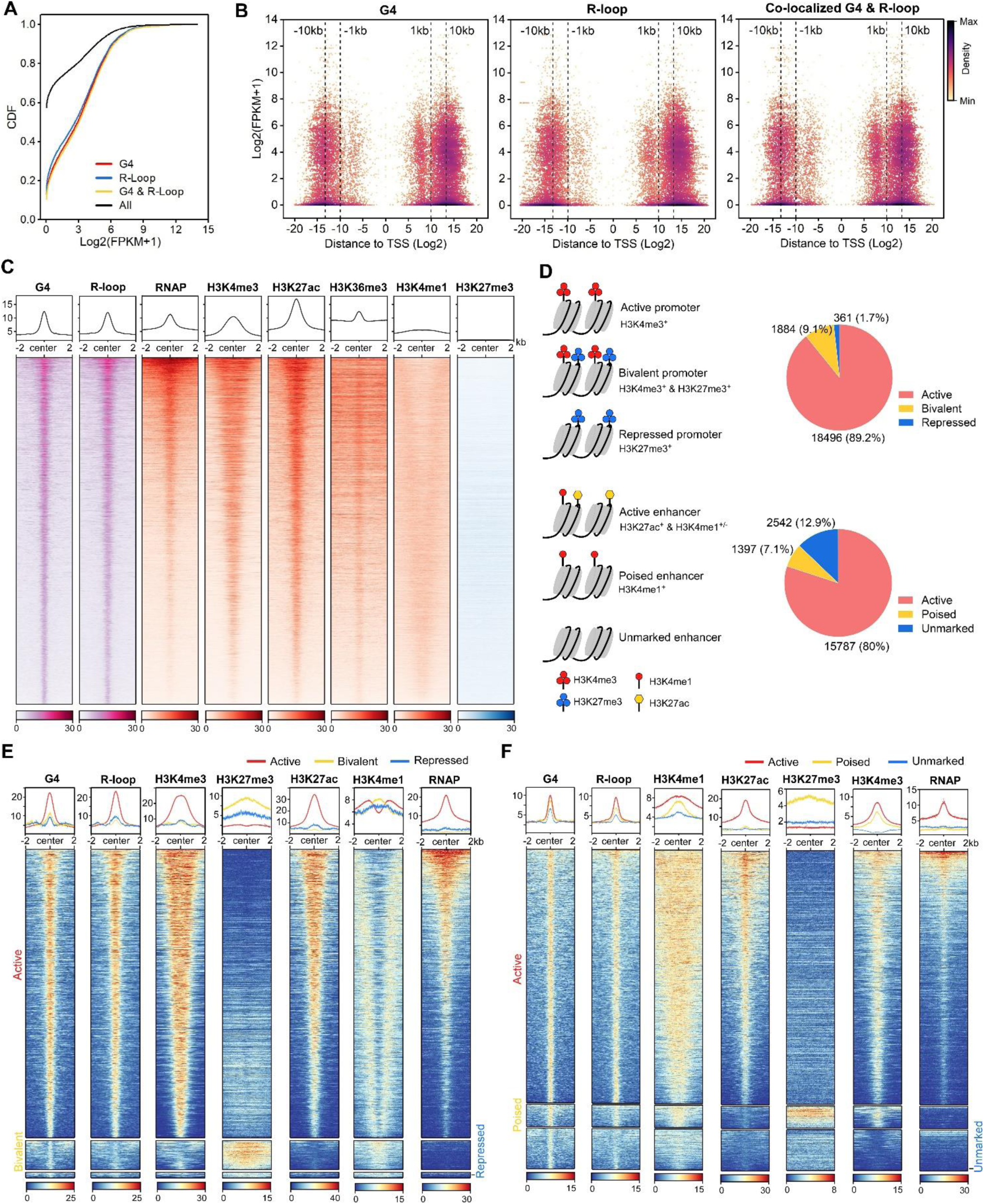
The co-localized G4s & R-loops are mainly localized in active promoters and enhancers. **A**, Cumulative distribution plot showing comparisons of FPKMs of G4-, R-loop-, and co-localized G4 & R-loop-associated genes and all genes in mESCs. **B**, Scatter plot showing the distributions of FPKMs of G4-, R-loop-, and co-localized G4 & R-loop -associated genes versus the distances of these peaks to nearest TSS. The distance is in bp. Color scales represent the density of dots. **C**, Heatmap showing the signal of representative HepG4-seq (G4), HBD-seq (R-loop), RNA polymerase II Ser5P, H3K4me3, H3K27ac, H3K36me3, H3K4me1, and H3K27me3 ±2kb around the center of the co-localized G4s & R-loops in mESCs. Color scales represent the density of the signals. The average signal is plotted at the top of each heatmap panel. **D**, Schematic of different types of promoters and enhancers. Pie chart showing the proportion of different types of promoters or enhancers that harbor the co-localized G4s & R-loops. **E-F**, Heatmap showing the signal of representative HepG4-seq (G4), HBD-seq (R-loop), RNA polymerase II Ser5P, H3K4me3, H3K27ac, H3K36me3, H3K4me1, and H3K27me3 ±2kb around the center of the co-localized G4s & R-loops in the different types of promoters (**E**) or enhancers (**F**) in mESCs. Color scales represent the density of the signals. The average signal is plotted at the top of each heatmap panel.

Extensive studies define promoters into active, bivalent and repressed states based on patterns of H3K4me3 and H3K27me3 (Fig. 5D); enhancers are defined as active, poised and unmarked states based on patterns of H3K27ac and H3K4me1 (Fig. 5D) (Atlasi and Stunnenberg, 2017; Bernstein et al., 2006; Bibikova et al., 2008; Calo and Wysocka, 2013; Heintzman et al., 2007). Notably, 20,741 and 19,726 co-localized G4s and R-loops are found in promoters and enhancers, respectively; 18,496 peaks are seen in active promoters; 15,787 peaks are seen in active enhancers (Fig. 5D, Supplementary Table 2). The co-localized G4s and R-loops in active promoters show high and almost equal signals of G4s and R-loops, and enrich H3K4me3, H3K27ac and RNAP (Fig. 5E). The co-localization of G4s, R-loops and RNAP at active promoters suggests that co-localized G4s and R-loops are likely linked to promoter-associated nascent RNAs (Core et al., 2008; Li and Fu, 2019; Preker et al., 2008; Seila et al., 2008). Interestingly, a medium level of H3K4me1 is present in a bimodal pattern beside co-localized G4s and R-loops at active promoters (Fig. 5E). The co-localized G4s and R-loops in bivalent promoters exhibit low signals of both G4s and R-loops, and overlap with a high level of H3K27me3, a medium level of H3K4me1, and a low level of H3K4m3 (Fig. 5E). The co-localized G4s and R-loops in active enhancers exhibit sharp peaks and well overlap with H3K27ac, H3K4me1, H3K4me3 and RNAP; the co-localized G4s and R-loops in poised enhancers enrich the active histone marks H3K4me1 and H3K4me3, and the repressive mark H3K27me3 while the signal of RNAP is very low (Fig. 5F). The unmarked enhancers-associated co-localized G4s and R-loops show low levels of all marks tested. Given that enhancer RNAs (eRNAs) have been widely identified as non-coding RNAs in enhancers and are functionally important for enhancer activity (Andersson et al., 2014; Kim et al., 2010; Sigova et al., 2015), the co-occupacy of eRNAs are likely involved in the formation of co-localized G4s and R-loops in enhancers and shed light on the new regulatory mechanism of eRNA action.

### Modulation of co-localized G4s and R-loops by the helicase Dhx9

Dhx9 (also known as RNA Helicase A) is a versatile helicase capable of directly resolving R-loops and G4s or promoting R-loop formation by unwinding secondary structures in the nascent RNA strand (Chakraborty and Grosse, 2011; Chakraborty et al., 2018; Cristini et al., 2018; Matsui et al., 2020; Tang et al., 2022; Yuan et al., 2021), and has been reported to play roles in DNA replication, transcription, translation, RNA processing and transport and maintenance of genomic stability (Aktas et al., 2017; Aratani et al., 2001; Chellini et al., 2022; Jain et al., 2013; Tang et al., 2022). Thus, Dhx9 is a promising regulator of co-localized G4s and R-loops.

To investigate the role of Dhx9 in modulating co-localized G4s and R-loops, we generated the Dhx9 knockout mESCs (*dhx9^KO^*) by CRISPR/Cas9-medidated gene editing (Ran et al., 2013). The depletion of Dhx9 in the *dhx9^KO^* mESC clone was confirmed by western blot assay (Fig. 6A) and immunofluorescence staining (Fig. 6B). We next examined the G4 an R-loop levels in the *dhx9^KO^* mESCs by performing HepG4-seq and HBD-seq. Notably, compared to the wildtype mESCs, a large amount of G4s or R-loops within co-localized G4s and R-loops exhibited significantly up-regulated or down-regulated signals in the *dhx9^KO^* mESCs (Fig. 6C, Supplementary Fig. 3A, Supplementary Table 2), suggesting that Dhx9 could unwind or promote the formation of co-localized G4s and R-loops in mESCs. Interestingly, only small proportion of co-localized G4s and R-loops displayed differential G4s and R-loops at the same time in the *dhx9^KO^*mESCs (Fig. 6D, Supplementary Fig. 3B), suggesting that Dhx9 cannot simultaneously unwind or promote G4s and R-loops within co-localized G4 and R-loop regions and that multiple helicases or regulators are required for modulating these regions.

**Fig. 6.**
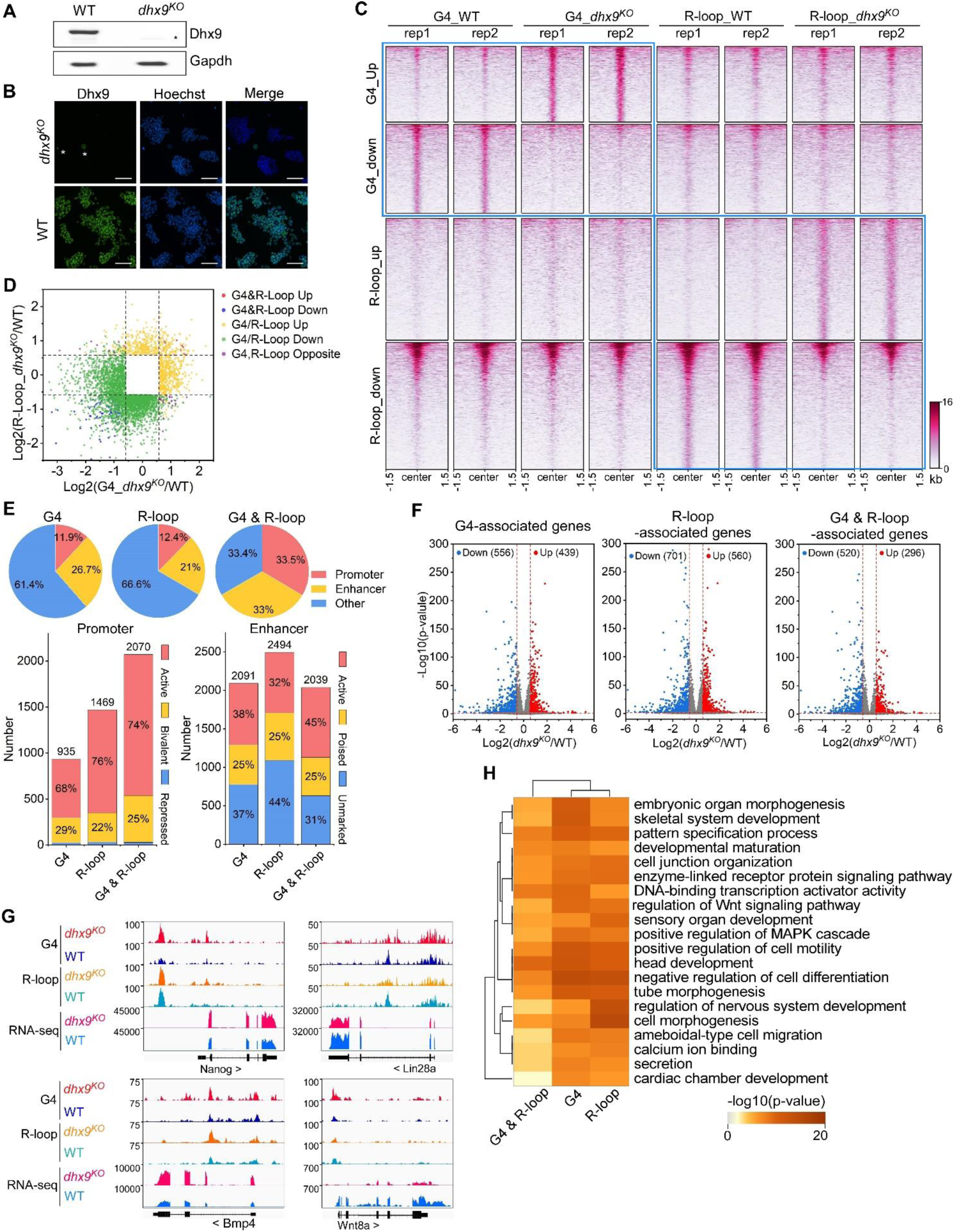
Modulation of the co-localized G4s & R-loops by the helicase Dhx9. **A**, Western blot showing the protein levels of Dhx9 and Gapdh in the wildtype (WT) and *dhx9^KO^* mESCs. The non-specific band is labeled with a star. **B**, Immunofluorescence staining of Dhx9 in the WT and *dhx9^KO^* mESCs cultured without MEFs feeder. Nuclei were stained with the Hoechst33342. The immunofluorescence signal of contaminated MEFs is labeled with a a star. Scale bar, 100 µm. **C**, Heatmap showing the signal of DNA G4 (HepG4-seq) and R-loop (HBD-seq) ±1.5kb around the center of significantly differential peaks (p-value < 0.05, fold change ≥ 1.5) in WT and *dhx9^KO^* mESCs. Two biologically independent replicates are shown. Color scales represent the density of the signals. **D**, Scatter plot showing distributions of foldchanges of differential G4s versus foldchanges of differential R-loop in WT and *dhx9^KO^* mESCs. “G4&R-loop Up”, both G4 and R-loop up-regulated; “G4&R-loop Down”, both G4 and R-loop down-regulated; “G4/R-loop Up”, G4 or R-loop up-regulated; “G4/R-loop Down”, G4 or R-loop down-regulated; “G4, R-loop Opposite”, G4 up-regulated and R-loop down-regulated, or, G4 down-regulated and R-loop up-regulated. **E**, Top: pie chart showing proportions of differential G4s, R-loops and co-localized G4s & R-loops in *dhx9^KO^* mESCs in promoters, enhancers and other regions; Bottom: bar chart showing numbers of differential G4, R-loops and co-localized G4s & R-loops in *dhx9^KO^* mESCs in different types of promoters or enhancers. **F**, Volcano plot showing distributions of G4-, R-loop-, or co-localized G4 & R-loop-associated genes differentially expressed in WT and *dhx9^KO^* mESCs. Significantly up-regulated (p-value < 0.05, fold change ≥ 1.5) and down-regulated (p-value < 0.05, fold change ≤ 0.67) genes in the *dhx9^KO^* mESCs are labeled with red and blue dots respectively. The numbers of up- or down-regulated genes are labeled on the plot. **G**, Representative genome browser tracks showing the G4 (HepG4), R-loop (HBD-seq), and RNA-seq signals in WT and *dhx9^KO^* mESCs along the indicated genomic loci. **H**, GO-based enrichment terms of G4-, R-loop-, or co-localized G4 & R-loop-associated genes differentially expressed in WT and *dhx9^KO^* mESCs were hierarchically clustered into a tree based on Kappa-statistical similarities among their gene memberships. The heatmap cells are colored by their p-values.

Given that co-localized G4s and R-loops have been shown to be enriched in active promoters and enhancers (Fig. 5D-F), the loss-of-Dhx9-induced differential co-localized G4s and R-loops preferentially localize in the active and bivalent promoters and all three types of enhancers (Fig. 6E). To explore the effect of Dhx9 on the transcription of co-localized G4s and R-loops -associated genes, we performed the RNA-seq on wild-type and *dhx9^KO^* mESCs. Differential gene expression analysis revealed 1647 significantly up-regulated genes and 1916 significantly down-regulated genes in absence of Dhx9 (Supplementary Fig. 3C-D). Importantly, loss of Dhx9 resulted in hundreds of G4-, R-loop-, and G4&R-loop-associated genes with significantly differential expression (Fig. 6F), suggesting that Dhx9 could regulate transcription by modulating G4s, R-loops and co-localized G4s and R-loops. Representative Dhx9-regulated locus are shown in Fig. 6G. GO analysis showed that co-localized G4s and R-loops-associated genes that showed differential expression after knocking out Dhx9 are mainly involved in negative regulation of cell differentiation, head development, positive regulation of cell motility, DNA-binding transcription activator activity, pattern specification process, embryonic organ morphogenesis, et al., suggesting that Dhx9 may regulate the cell fate of mESCs by modulating co-localized G4s and R-loops (Fig. 6H). Coinciding with GO analysis, Dhx9 knockout in the mouse causes embryonic lethality and Dhx9 knockdown leads to large structural changes in chromatin and eventually cell death (He et al., 2008; Zhang et al., 2004). Heterozygous loss-of-function variants of DHX9 are associated with neurodevelopmental disorders in human (Yamada et al., 2023).

### Characterization of co-localized G4s and R-loops directly bound by Dhx9

Tens of helicases or regulators have been reported to directly resolve or stabilize G4s or R-loops (Mendoza et al., 2016; Varshney et al., 2020; Yang et al., 2023). Interestingly, loss of Dhx9 caused 30 of these helicases/regulators to be significantly differentially expressed (Fig. 7A) and Dhx9 physically interacts with at least ten of them based on the STRING protein-protein interaction network database (Szklarczyk et al., 2023) (Supplementary Fig. 3E). These data suggest that Dhx9 could also indirectly modulate G4s and R-loops by affecting other helicases or regulators. Thus, to explore the direct target co-localized G4s and R-loops of Dhx9, we performed the CUT&Tag-seq using the Dhx9 antibody (Kaya-Okur et al., 2019) and revealed 54,982 Dhx9 binding peaks in wild-type mESCs (Fig. 7B). Notably, 65.5% Dhx9 binding peaks well overlapped with 69.9% co-localized G4s and R-loops in mESCs (Fig. 7C-D), suggesting that Dhx9 is a direct and major regulator of co-localized G4s and R-loops in mESCs. Motif analysis showed that G-rich sequences are highly enriched in the Dhx9 binding peaks overlapping with co-localized G4s and R-loops in mESCs (Fig. 7E), further demonstrating that Dhx9 directly bind to co-localized G4s and R-loops that harbor G-rich sequences as shown in Fig. 4G.

**Fig. 7.**
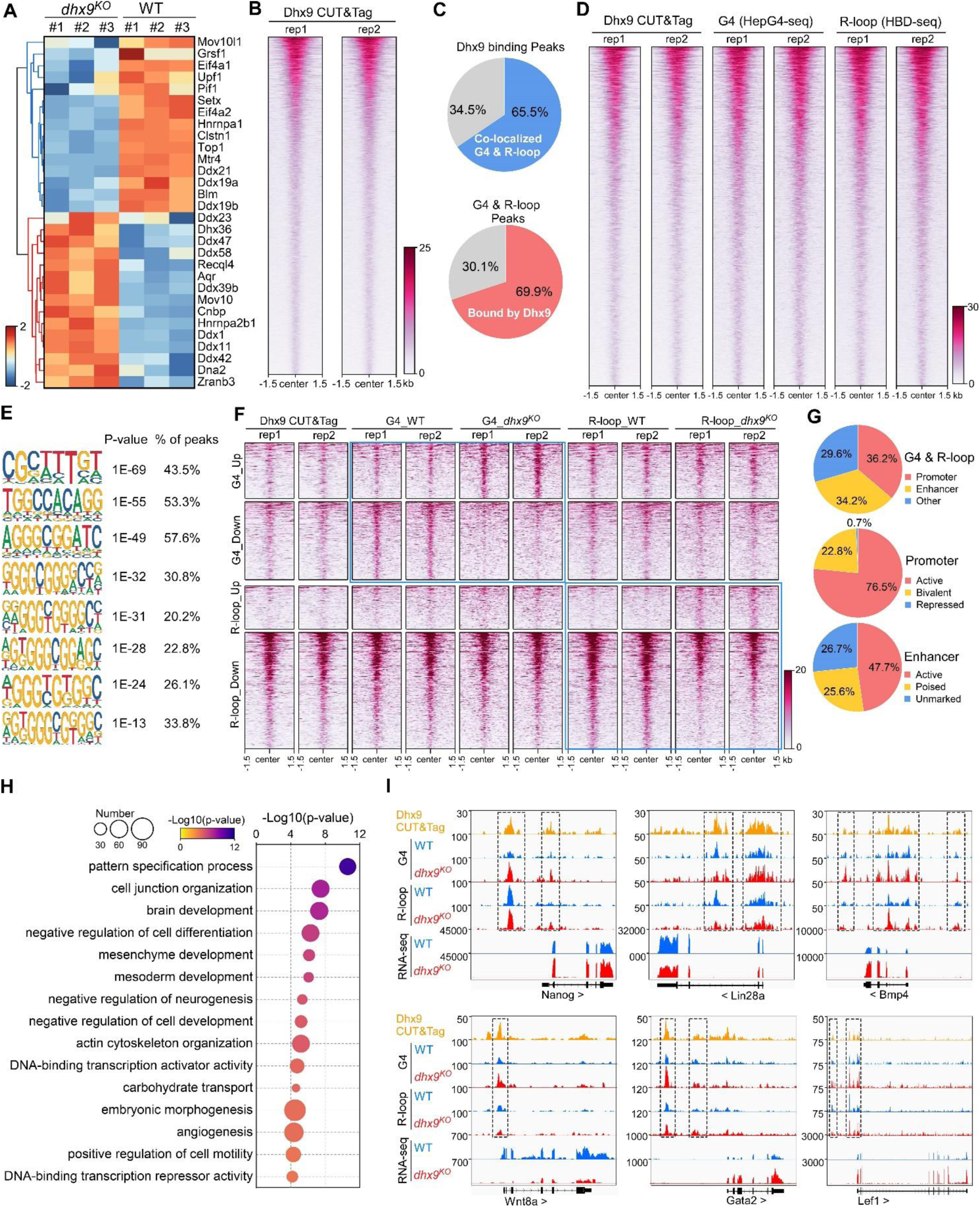
Characterization of the co-localized G4s & R-loops directly bound by Dhx9. **A**, Heatmap showing expression levels of resolving helicases or regulators of G4s and/or R-loops differentially expressed in WT and *dhx9^KO^* mESCs. Color scales represent the normalized expression levels. **B**, Heatmap showing the signal of Dhx9 CUT&Tag reads ±1.5kb around the center of peaks in WT mESCs. Two biologically independent replicates are shown. Color scales represent the density of the signals. **C**, Pie chart showing proportions of the overlapping peaks between Dhx9 binding peaks and the co-localized G4 & R-loops peaks. **D**, Heatmap showing the signal of Dhx9 CUT&Tag, HepG4-seq and HBD-seq ±1.5kb around the center of Dhx9-bound Co-localized G4 & R-loop peaks in mESCs. Two biologically independent replicates are shown. Color scales represent the density of the signals. **E**, The top enriched motifs of Dhx9 binding peaks overlapping with the co-localized G4 & R-loops in mESCs. **F**, Heatmap showing the signal of Dhx9 CUT&Tag, HepG4-seq and HBD-seq ±1.5kb around the center of Dhx9-bound significantly differential Co-localized G4 & R-loop peaks (p-value < 0.05, fold change ≥ 1.5) in WT and *dhx9^KO^* mESCs. Two biologically independent replicates are shown. Color scales represent the density of the signals. **G**, Pie chart showing proportions of differential Dhx9-bound co-localized G4 & R-loops in *dhx9^KO^* mESCs in different types of promoters or enhancers. **H**, Top enriched GO terms in Dhx9-bound co-localized G4 & R-loop-associated genes that are differentially expressed in WT and *dhx9^KO^* mESCs. The bubble size represents the number of genes in each indicated term. The color scale represents the p-value. **I**, Representative genome browser tracks showing the Dhx9 CUT&Tag, G4 (HepG4), R-loop (HBD-seq), and RNA-seq signals in WT and *dhx9^KO^* mESCs along the indicated genomic loci. The dashed box highlights Dhx9-bound significantly differential co-localized G4 & R-loop peaks in *dhx9^KO^* mESCs.

We next compared the Dhx9-bound co-localized G4s and R-loops in wild-type and *dhx9^KO^* mESCs, and identified 1,382 significantly up-regulated peaks (823 with increased G4s and 559 with increased R-loops) and 4,789 significantly down-regulated peaks (2,278 with decreased G4s and 2,511 with decreased R-loops) (Fig. 7F, Supplementary Fig. 3F), accounting for ∼50-75% of differential co-localized G4s and R-loops in absence of Dhx9 (Fig. 6C). Analysis of the genomic distribution of Dhx9-bound differential co-localized G4s and R-loops found that these peaks are mainly localize in the active and bivalent promoters and all three types of enhancers (Fig. 7G). The Dhx9-bound differential co-localized G4s and R-loops were linked to 852 genes with significantly differential expression (Supplementary Fig. 3G, Supplementary table 2), which are enriched in GO terms related to pattern specification, cell junction organization, brain development, negative regulation of cell differentiation, mesenchyme/mesoderm development, embryonic morphogenesis, et al. (Fig. 7H). Several key regulators of mouse embryonic stem cell and embryonic development, such as Nanog, Lin28a, Bmp4, Wnt8a, Gata2, and Lef1, were shown to be transcriptionally regulated by Dhx9 through direct modulation of their associated G4s and R-loops (Fig. 7I). These data suggest that Dhx9 significantly contributes to transcriptional regulation of co-localized G4s and R-loops -associated genes.

### Dhx9 regulates the cell fate of mESCs

To understand the role of Dhx9 in regulating the cell fate of mESCs, we first examined the RNA levels of several key genes that maintain the pluripotency of mESCs by quantitative RT-PCR (qRT-PCR) and found that the RNA levels of Lin28a and Oct4 were significantly decreased and the RNA level of Nanog was significantly increased when Dhx9 was knocked out (Fig. 8A). Western blot assay showed that *dhx9^KO^* mESCs produce obviously lower level of Lin28a protein than wild-type mESCs, consistent with its RNA level, whereas in contrast to the RNA level, the protein level of Nanog was significantly decreased in *dhx9^KO^* mESCs, suggesting that Dhx9 directly or indirectly modulates the translation of Nanog (Fig. 8B). In line with the western blot assay, immunofluorescence staining of WT and *dhx9^KO^*mESCs showed that the loss of Dhx9 leads to reduced protein level of Nanog, but not Oct4, while *dhx9^KO^* mESCs exhibited normal morphology (Fig. 8C). The mESCs can be maintained in a proliferative state for prolonged periods, which was known as “self-renewal” (Liang and Zhang, 2013; Murry and Keller, 2008). Nanog and Lin28a have been reported to promote embryonic stem self-renewal (Chambers et al., 2003; Mitsui et al., 2003; Xu et al., 2009). Coinciding with reduced levels of Nanog and Lin28a proteins, *dhx9^KO^* mESCs was shown to modestly arrest at S phase of the cell-cycle (Fig. 8D) and exhibited significantly attenuated proliferation capacity (Fig. 8E), suggesting that Dhx9 regulates the self-renewal of mESCs.

**Fig. 8.**
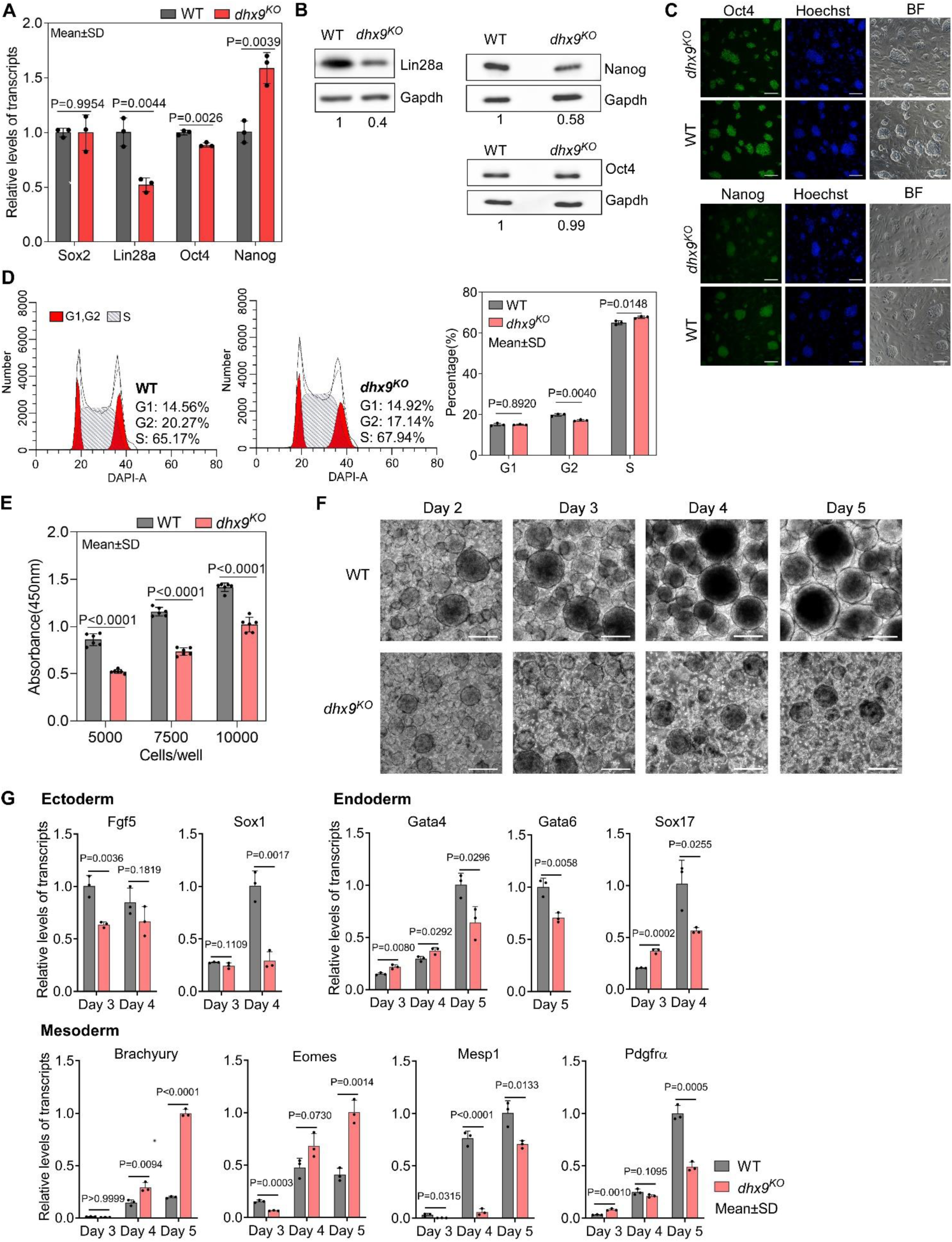
Dhx9 regulates the cell fate of mouse embryonic stem cells. **A**, Relative RNA levels of indicated genes in the WT and *dhx9^KO^* mESCs that were measured by quantitative RT-PCR (qRT-PCR). Data are means ± SD; n=3, significance was determined using the two-tailed Student’s *t*-test. **B**, Western blot showing the protein levels of indicated genes in the WT and *dhx9^KO^* mESCs. The normalized relative protein levels were labeled below each panel, where gel images were quantified by ImageJ and the level of Gapdh was used for normalization. **C**, Immunofluorescence staining of Oct4 and Nanog in the WT and *dhx9^KO^* mESCs cultured on MEF feeder. Nuclei were stained with the Hoechst33342. BF, bright field. Scale bar, 100 µm. **D**, Cell cycle profiles determined by flow cytometry of DAPI staining in the WT and *dhx9^KO^* mESCs. The proportions of different phases of cell cycle were analyzed by ModFit. Data are means ± SD; n=3, significance was determined using the two-tailed Student’s *t*-test. **E**, Cell proliferation rate of WT and *dhx9^KO^* mESCs were measured by CCK-8 cell proliferation assay. The number of cells seeded at the beginning was labeled at the x-axis. Absorbance at 450 nm was determined after 2 days of culture. Data are means ± SD; n=6, significance was determined using the two-tailed Student’s *t*-test. **F**, Pictures of embryoid bodies at indicated days of *in vitro* differentiation of WT and *dhx9^KO^* mESCs. Scale bar, 200 µm. **G**, Relative RNA levels of indicated genes in the WT and *dhx9^KO^* embryoid bodies at indicate days of *in vitro* differentiation that were measured by qRT-PCR. Data are means ± SD; n=3, significance was determined using the two-tailed Student’s *t*-test.

mESCs are pluripotent stem cells which are able to differentiate into three germ lineages (Murry and Keller, 2008; Young, 2011). In the absence of differentiation inhibitor LIF, mESCs cultured in suspension spontaneously form three-dimensional aggregates called embryoid bodies (EBs), which could recapitulate many aspects of early embryogenesis, including the induction of three early germ lineages (Simunovic and Brivanlou, 2017). To understand role of Dhx9 in regulating the pluripotency of mESCs, we performed the EB assay using the wild-type and *dhx9^KO^* mESCs. As the EB differentiation progressed, loss of Dhx9 resulted in apparently smaller and fewer EBs than wild-type cells (Fig. 8F). At the same time, we collected EBs at different days of EB differentiation and examined the RNA levels of well-known markers of three germ lineages by qRT-PCR. As shown in Fig. 8G, all maker genes tested displayed significantly differential expression, suggesting that Dhx9 regulates the pluripotency of mESCs, which is in line with the GO enrichment results of Dhx9-regulated co-localized G4s and R-loops-associated genes in Fig. 7H. Taken together, Dhx9 regulates the self-renewal and differentiation capacities of mESCs.

### Comparisons of HepG4-seq and HBD-seq with previous methods

To compare the performance of HepG4-seq and BG4 CUT&Tag, we analyzed the data using the same bioinformatic pipeline. As shown in Fig.9A, 80% (5,459) and 71% (67,935) HepG4-seq peaks overlap with BG4 CUT&Tag peaks in HEK293 cells and mESCs, respectively. SEACR is highly selective peak caller for CUT&Tag and CUT&Run (Meers et al., 2019). With the help pf SEACR, BG4 CUT&Tag obtained a much larger number of peaks in HEK293 cells and mESCs than previously reported MACS2-identified peaks. Surprisingly, 71,363 and 103,303 consensus peaks were identified by BG4 CUT&Tag alone in HEK293 cells and mESCs, respectively (Fig.9A), suggesting that BG4 antibody may promote the folding of G4 sequences during BG4-seq or that BG4 antibody is more sensitive than G4-hemin-induced proximal biotinylation in recognizing native G4s. As shown in Fig.9B, the patterns of BG4 CUT&Tag peaks overlap well with those of PQS while HepG4-seq peaks exhibit a much sharper shape and better resolution around the center of PQS patterns, indicating that BG4 may promote the folding of G4 motifs. Interestingly, the signal intensities of the HepG4-seq peaks are significantly lower than those of BG4 CUT&Tag in HEK293 cells, but obviously higher than those of BG4 CUT&Tag in mESCs. In the *in vitro* hemin-G4 induced self-biotinylation assay, parallel G4s exhibit higher peroxidase activities than anti-parallel G4s (Lat et al., 2020). The signal intensities changes of HepG4-seq in different cells possibly reflect the dynamics of G4 conformation. In the future, people may need to combine HepG4-seq and BG4s-eq to carefully explain the endogenous G4s.

**Fig. 9.**
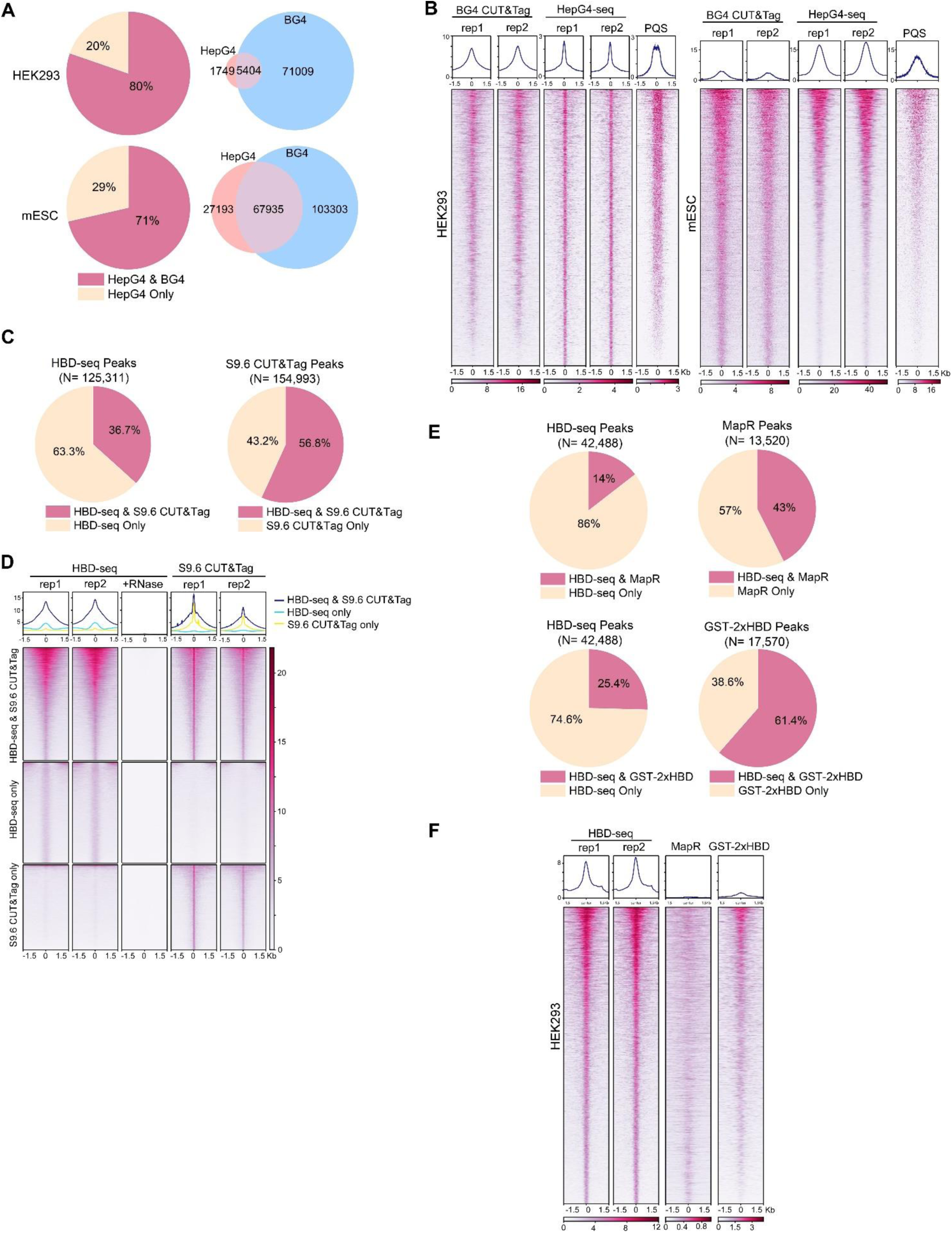
Comparisons of HepG4-seq and HBD-seq with previous methods. **A**, Pie charts showing the percentages of HepG4-seq peaks overlapping with peaks identified by BG4 CUT&Tag; Venn diagrams comparing HepG4-seq peaks and BG4 CUT&Tag seqs. **B**, Heatmap showing the signal of BG4 CUT&Tag, HepG4-seq and maxScores of PQS ±1.5kb around the center of peaks identified by both BG4 CUT&Tag and HepG4-seq in HEK293 cells or mESCs. Color scales represent the density of the signals. Top: Profile plot showing the average signal. Rep1/rep2, two biologically independent replicates. **C**, Pie charts showing the overlapping percentages of peaks between HBD-seq and S9.6 CUT&Tag in mESCs. **D**, Heatmap showing the signal of HBD-seq and S9.6 CUT&Tag ±1.5kb around the center of peaks identified by HBD-seq and S9.6 CUT&Tag in mESCs. Color scales represent the density of the signals. Top: Profile plot showing the average signal. Rep1/rep2, two biologically independent replicates. **E**, Pie charts showing the overlapping percentages of peaks between HBD-seq and MapR or GST-2xHBD CUT&Tag in HEK293 cells. **F**, Heatmap showing the signal of HBD-seq, MapR and GST-2xHBD CUT&Tag ±1.5kb around the center of peaks identified by HBD-seq in HEK293 cells. Color scales represent the density of the signals. Top: Profile plot showing the average signal. Rep1/rep2, two biologically independent replicates.

During the past decade, R-loops were mapped using either S9.6 monoclonal antibody or catalytically inactive ribonuclease H1 for specific DNA-RNA hybrid. First, we compared the HBD-seq with S9.6 CUT&Tag in mESCs using the same bioinformatic pipeline and SEACR for peak calling. As a result, HBD-seq and S9.6 CUT&Tag identified similar numbers of peaks (125,311 for HBD-seq, 154,993 for S9.6 CUT&Tag), and 45,989 HBD-seq peaks overlap with 88,036 S9.6 peaks, indicating that S9.6 peaks are possibly narrower than HBD-seq peaks. As shown in Fig.9D, the overlapping peaks showed similar heatmap patterns and signal intensities between HBD-seq and S9.6 CUT&Tag while, notably, peaks of S9.6 CUT&Tag exhibit a much sharper signals at the center of peaks, which reflect the difference of binding modules of HBD and S9.6. GST-2xHBD and S9.6 mediated CUT&Tag were shown to generate highly similar native R-loop profiles (Wang et al., 2021). HBD-seq, as an optimized version of GST-2xHBD CUT&Tag, still generated ∼80,000 extra peaks compared to S9.6 CUT&Tag, suggesting that the status of the recombinant 2xHBD protein could substantially affect R-loop mapping. To understand the difference among inactive ribonuclease H1-based methods, we compared the HBD-seq, MapR (inactive ribonuclease H1-mediated CUT&Run) (Yan et al., 2019) and GST-2xHBD CUT&Tag (Wang et al., 2021) and re-analyzed the data using the same pipeline and peak caller SEACR. In HEK293 cells, HBD-seq, MapR, and GST-2xHBD CUT&Tag identified 42,488, 12,520, and 17,570 peaks respectively; 5,752 MaR peaks and 11,701 GST-2xHBD CUT&Tag peaks overlap with HBD-seq peaks (Fig.9E). Notably, the heatmap of all HBD-seq peaks showed an obviously stronger signal intensities and sharper peak shapes than MapR and GST-2xHBD CUT&Tag, suggesting that HBD-seq provides superior R-loop signals. In consideration of the difference in DNA-RNA hybrid binding affinity and/or specificity between HBD and S9.6, the HBD-seq and S9.6 CUT&Tag may need to be combined together to carefully explain the native R-loops.

## Discussion

In this study, we developed the new method “HepG4-seq” and optimized the RNase H1 HBD domain-based HBD-seq to robustly map endogenous G4s and R-loops, respectively, in living cells with high specificity. Using the HepG4-seq and HBD-seq, we systematically characterized the native co-localized G4s and R-loops in HEK293 cells and mESCs, and revealed that co-localized G4s and R-loops are dynamically altered in a cell type-dependent way and largely localized at active promoters and enhancers of transcriptional active genes. Small molecules-induced inhibition of helicases BLM or WRN resulted in significant accumulation of G4s within co-localized G4s and R-loops and at the same time leaded to genes with significantly differential expression in HEK293 cells that are enriched in the processes related to cell cycle, DNA metabolic, DNA damage response, chromatin binding, et al. Furthermore, we characterized the helicase Dhx9 as a key regulator of co-localized G4s and R-loops which efficiently unwinds or promotes co-localized G4s and R-loops, and illustrated that depletion of Dhx9 significantly altered the transcription of co-localized G4s and R-loops-associated genes that are enriched in embryonic development, cell differentiation and germ lineage development, et al. Therefore, loss of Dhx9 apparently impaired the self-renewal and pluripotency of mESCs.

In this study, we utilized a low dosage of hemin, similar to the physiological concentration in normal human erythrocytes, to spark the peroxidase activity of endogenous G4s without significantly altering the levels of native G4s (Supplementary Fig. 1B-C) and further robustly biotinylated G4s themselves by G4-hermin complex-mediated proximity labeling in just one minute in living cells (Cheng et al., 2009; Einarson and Sen, 2017; Lat et al., 2020; Li et al., 2016; Stadlbauer et al., 2021; Yang et al., 2011). In consideration of the high affinity and specificity, the recombinant streptavidin monomer (Lim et al., 2013) is able to recognize the biotinylated G4s with high sensitivity and specificity and thereby yield robust CUT&Tag signals with the help of Moon-tag system (Boersma et al., 2019). Therefore, our HepG4-seq strategy is able to robustly and specifically capture native G4s. In HEK293 cells, HepG4-seq uncovered 6,799 consensus G4 peaks under wild-type status and 77,003 G4 peaks in the present of BLM/WRN inhibitors, suggesting that HepG4-seq is capable of detecting endogenous G4s with high sensitivity. Yet, the hemin-G4 complex induced peroxidase activities were reported to be variable between parallelled and anti-parallelled G4s in the *in vitro* assay (Lat et al., 2020), although only 2-4 synthesized short DNA fragments were used for the test. Given that a method of genome-widely distinguishing parallelled and anti-parallelled G4s is lacking, HepG4-seq and BG4-seq may be combined to characterize the conformation of G4s.

Notably, we also discovered that the native co-localized G4s and R-loops landscape is altered in a cell-dependent manner and that approximately 10 folds more peaks were observed in mESCs than in HEK293 cells, which reflects that co-localized G4s and R-loops have the potential to regulate the complex pluripotency network in mESCs. CTCF is a key regulator of genome organization and gene expression (Ong and Corces, 2014). Recently, Wulfridge et al. reported that CTCF-bound regions are enriched for both R-loops and G4s and G4s associated with R-loops promote CTCF binding (Wulfridge et al., 2023). Interestingly, the enriched motif with the most significant p-value in mESCs co-localized G4s and R-loops (Fig. 4G) well matches the motif of CTCF ChIP-seq, suggesting that co-localized G4s and R-loops may be able to modulate the CTCF binding. To be noticed, the co-localized G4s and R-loops were identified by bioinformatically intersection analysis. The heterogenous distribution between cells will give false positive co-occurrence patterns. Recently, multi-CUT&Tag, a method for simultaneous mapping of multiple chromatin proteins, has been reported (Bartosovic and Castelo-Branco, 2023; Gopalan et al., 2021; Meers et al., 2023). As the multi-CUT&Tag, we could combine the HepG4 and HBD-seq to simultaneously map the co-localized G4s and R-loops even in the single cell level.

Furthermore, while 47,857 co-localized G4s and R-loops are directly bound by Dhx9 in the wild-type mESCs (Supplementary Table 2), only 4,060 of them display significantly differential signals in absence of Dhx9. In addition, a limited number of co-localized G4s and R-loops showed differential G4s and R-loops at the same time in the *dhx9^KO^* mESCs. These data suggest that redundant regulators exist. It is worth noting that depletion of Dhx9 significantly altered the transcription of 30 known G4s and/or R-loops helicases/regulators (Fig. 7A) and that half of these helicases/regulators are able to establish physical interaction network (Supplementary Fig. 3E). Due to the redundance and compensatory roles of G4 and R-loop regulators, a comprehensively mechanistic study of G4 and R-loop interplay in living cells is difficult and lacking in the field. A degron system-mediated simultaneous and/or stepwise degradation system of multiple regulators will help us elucidate the interplaying effects between G4s and R-loops.

Taken together, our study provides new insights into exploring regulatory roles of co-localized G4s and R-loops in development and disease.

## Data availability

The HepG4-seq, HBD-seq, BG4-seq, Dhx9 CUT&Tag and RNA-seq data have been deposited to the Gene Expression Omnibus (accession code GSE254764 and GSE254763). The ChIP-seq data of histone markers and RNAP are openly available in GNomEx database (accession number 44R) (Wamstad et al., 2012).

## Acknowledgements

Z.X. is supported by the National Key Research and Development Program of China, Stem Cell and Translational Research (2018YFA0109200) and National Natural Science Foundation of China (General Program No. 31970600).

## Author information

T.L.: study design, experimental work, data interpretation, and writing of the manuscript. X.S.: study design, experimental work, data interpretation, and writing of the manuscript. Y.R.: bioinformatic analysis, data interpretation, and manuscript review. H.L.: experimental work, data interpretation. Y.L.: data review, manuscript review. C.C.: data review, manuscript review. L.Y.: study design, experimental work, data interpretation, writing and review of the manuscript. Z.X.: study design, data interpretation, bioinformatic evaluation, writing and review of the manuscript, supervision of the work.

---

**Supplementary Fig. 1.**
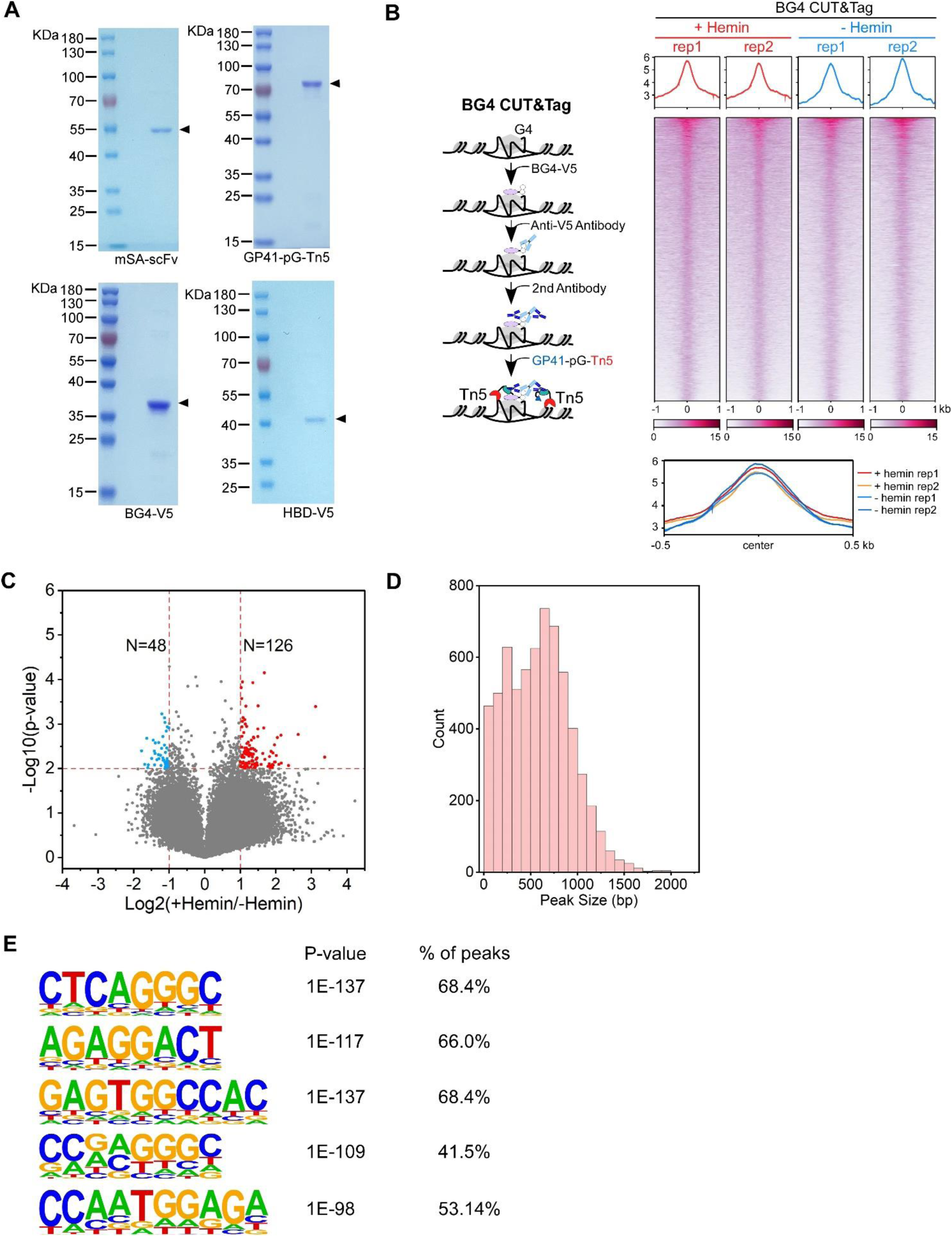
**A**, The Coomassie blue stained SDS-PAGE gel showing the purified recombinant mSA-scFV (mSA and anti-GP41 scFV), GP41-pG-Tn5 (GP41 tag, protein G and Tn5), BG4-V5 (G4 scFV BG4 and V5 tag), and HBD-V5 (N-terminal hybrid-binding domain of RNase H1, EGFP and V5 tag). **B**, Left: Schematic of the BG4 CUT&Tag procedure. Right top: Heatmap showing the signal of BG4 CUT&Tag ±1.5kb around the center of peaks in HEK293 cells treated with or without 25 µM hemin. Two biologically independent replicates are shown. Color scales represent the density of the signals. The average signal is plotted at the top of each heatmap panel. Right bottom: Profile plot showing the average signal of BG4 CUT&Tag reads around the center of peaks in HEK293 cells. **C**, Volcano plot showing distributions of differential BG4 CUT&Tag peaks in HEK293 cells treated with or without 25 µM hemin. Significantly up-regulated (p-value < 0.01, fold change ≥ 2) and down-regulated (p-value < 0.01, fold change ≤ 0.5) peaks are labeled with red and blue dots respectively. The numbers of up- or down-regulated peaks are labeled on the plot. **D**, Bar chart showing the distribution of co-localized G4 & R-loop peak sizes in HEK293 cells. **E**, The top enriched motifs of the extra accumulated peaks induced by inhibition of WRN and BLM in the figure 1 H-I.

**Supplementary Fig. 2.**
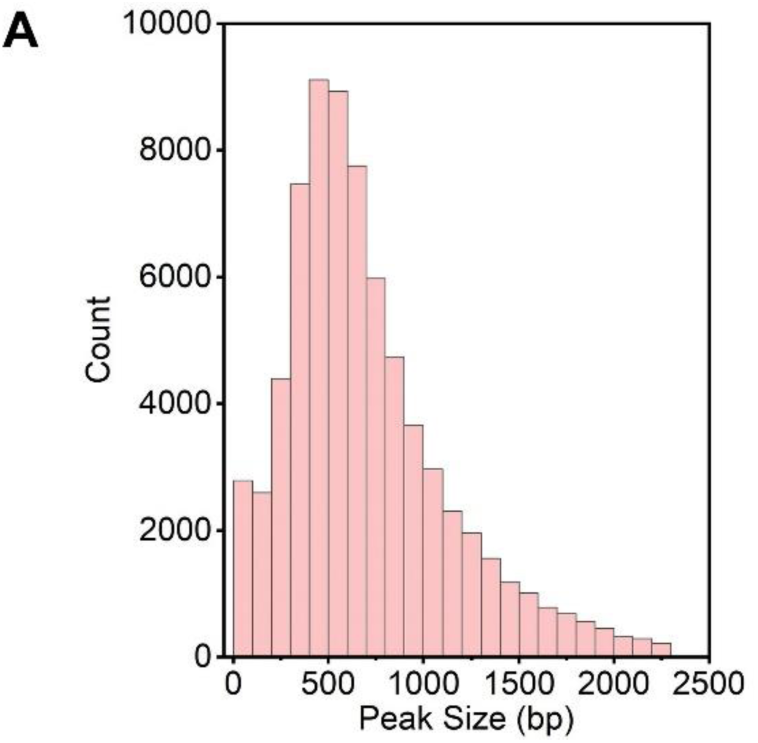
**A**, Bar chart showing the distribution of co-localized G4 & R-loop peak sizes in mESCs. Different, the strand of PQS localized is the same to the template strand of the nearest gene; NA, no PQS.

**Supplementary Fig. 3.**
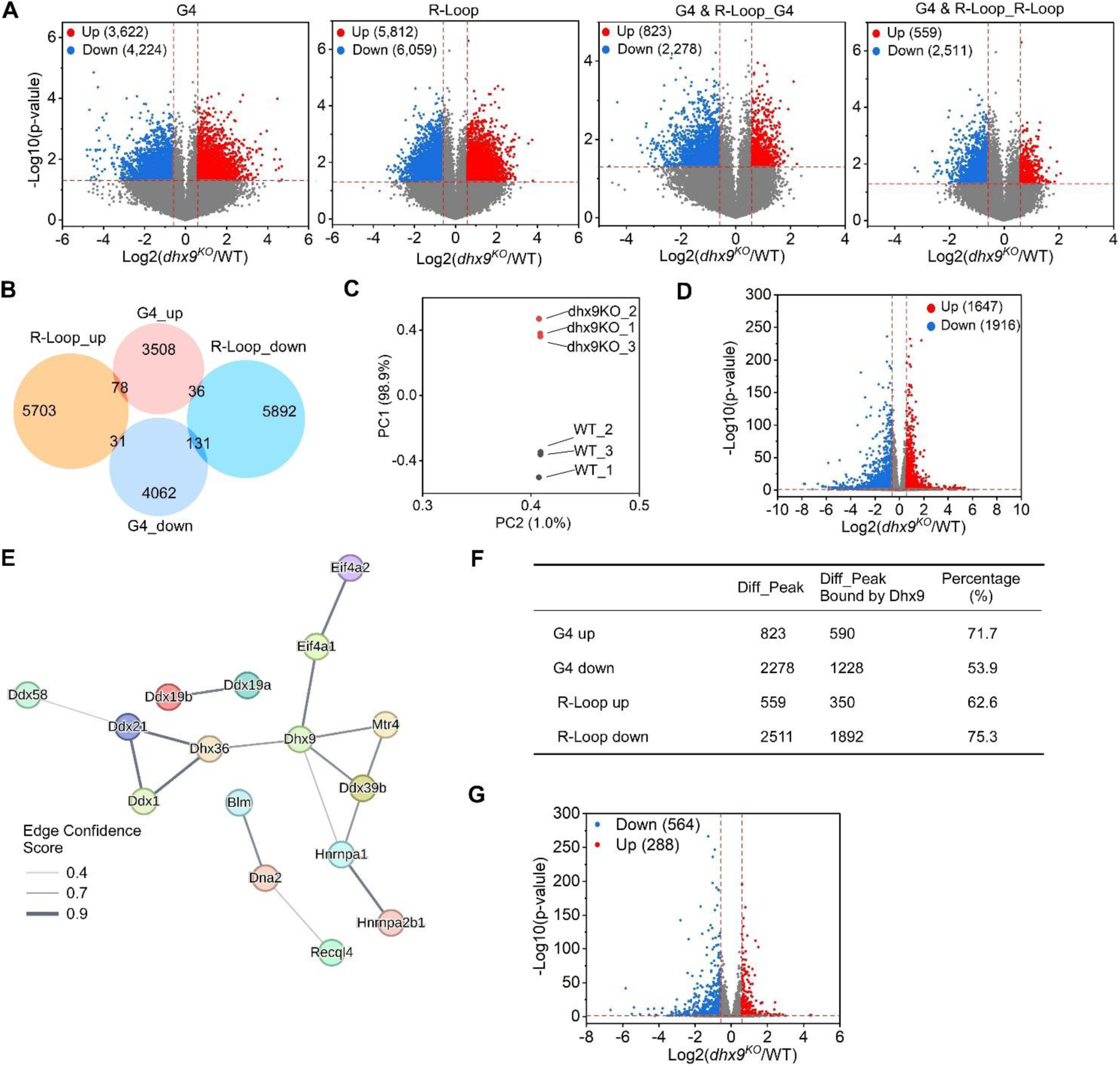
**A**, Volcano plot showing distributions of differential G4s and R-loops in WT and *dhx9^KO^* mESCs. Significantly up-regulated (p-value < 0.05, fold change ≥ 1.5) and down-regulated (p-value < 0.05, fold change ≤ 0.67) G4s or R-loops in the *dhx9^KO^* mESCs are labeled with red and blue dots respectively. The numbers of up- or down-regulated G4s or R-loops are labeled on the plot. **B**, Venn diagram comparing differential G4s or R-loops in WT and *dhx9^KO^* mESCs. **C**, Principal component analysis (PCA) of RNA levels of WT and *dhx9^KO^* mESCs. **D**, Volcano plot showing distributions of genes differentially expressed in WT and *dhx9^KO^* mESCs. Significantly up-regulated (p-value < 0.05, fold change ≥ 1.5) and down-regulated (p-value < 0.05, fold change ≤ 0.67) genes in the *dhx9^KO^* mESCs are labeled with red and blue dots respectively. The numbers of up- or down-regulated genes are labeled on the plot. **E**, The STRING protein-protein physical interaction network of G4s and/or R-loops resolving helicases or regulators that differentially expressed in WT and *dhx9^KO^* mESCs. The line thickness indicates the edge confidence scores that report the strength of data support from STRING database. The minimum required edge confidence score is 0.4. **F**, Table summarizing the differential co-localized G4s & R-loops in absence of Dhx9 and the differential co-localized G4s & R-loops directly bound by Dhx9. **G**, Volcano plot showing distributions of genes differentially expressed in WT and *dhx9^KO^* mESCs. Significantly up-regulated (p-value < 0.05, fold change ≥ 1.5) and down-regulated (p-value < 0.05, fold change ≤ 0.67) genes in the *dhx9^KO^* mESCs are labeled with red and blue dots respectively. The numbers of up- or down-regulated genes are labeled on the plot.

